# Using a 3-step semi-automated image analysis pipeline to quantify differences in VE-cadherin junction morphology in lymphatic endothelial cells in vitro

**DOI:** 10.64898/2026.10.05.756815

**Authors:** Stephen Larson, Annabelle Feller, Catherine Whittington

## Abstract

Lymphatic endothelial cell (LEC) function is strongly tied to transitions between button- and zipper-like junctions that control fluid flow into vessels. Published methods quantifying junction morphology are limited to single cell analysis or use few quantitative metrics to describe junctions across LEC monolayers. Therefore, a novel 3-step image analysis pipeline was developed to identify new metrics that quantify differences in button- and zipper-like junctions in LECs. The pipeline used Cellpose, CellProfiler, and Fiji to analyze VE-cadherin expression across lymphatic vessels within a 3D microfluidic chip and quantify VE-cadherin radial intensity, number of objects, object size, branchpoints, endpoints, and junction thickness. Tumor necrosis factor (TNF)-α exposure produced button-like junctions, and unexposed cells were zippered. Radial intensity was heightened at the cell membrane in zipper-like junctions while perinuclear intensity increased for button-like junctions. Increased branchpoints, endpoints, objects per cell, and junction thickness were associated with junction buttoning, while increased object size was associated with junction zippering. Principle component analysis results reinforced these trends and identified metric combinations that represented each junctional state. Overall, this semi-automated pipeline quantified differences in VE-cadherin junction morphology across LEC monolayers and generated new quantitative metrics to expand the tools available for studying junction morphology in lymphatic vasculature.

## 1. Introduction

Lymphatic vasculature is vital to maintain tissue homeostasis and facilitate adaptive immune response by draining interstitial fluid from tissues to lymph nodes[1], largely driven by tissue fluid moving passively along a pressure gradient controlled by cell junctions. The formation of cell junctions between lymphatic endothelial cells (LEC) impacts functional tissue properties such as permeability and overall vascular integrity[2]. Junction protein expression and junction morphology are key indicators of vascular function. Yet, assessments of junction morphology have been limited by the large degree of heterogeneity in junction morphology across monolayers, making automation for quantitative measurements a challenge[3]. Moreover, researchers are limited in their ability to correlate junction features with functional outcomes to better predict the dynamics of molecules and cells crossing lymphatic vessel walls under normal and diseased conditions.

Junction morphology for both tight junctions (e.g., ZO-1, occludin, claudin-5) and cadherin proteins (e.g., VE-cadherin, β-catenin, p120-catenin) are divided into either button-like or zipper-like morphologies that control fluid flow[1,2,4]. Button-like junctions are discontinuous junctions dotting the boundaries of lymphatic capillaries to allow fluid to enter lymphatic vasculature. This morphology is usually observed in healthy states. Zipper-like junctions are long continuous junctions that prevent fluid from crossing vessel walls and are characteristic of vessels upstream to the capillaries (i.e., collecting lymphatics). In pathological conditions (i.e. fibrosis), junction morphology can change from a button- to zipper-like morphology in lymphatic capillaries [5,6] which disrupts lymphatic function and can lead to edema and impaired immune response. Inflammatory molecules can act on junctions to sustain a zippered state, but some molecules (i.e., TNF-α) can induce junctions to revert to button-like, thus restoring lymphatic function to allow cells and molecules to cross the vessel wall[6]. Since junctional state is important for regulating lymphatic vascular function, assessments have often focused on measuring function or evaluating the expression of junction proteins rather than direct quantitative characterization of junction morphology.

Assays that directly measure lymphatic vascular function are often used to infer junction formation rather than measure it directly, and they often involve factors that are unrelated to junction morphology. Therefore, direct junction analysis is often paired with these assays to establish structure-function relationships. Permeability and transendothelial electrical resistance (TEER) are two examples of functional assays that are paired with junction analysis[7–12]. Permeability assays measure the transport of fluid or movement of cells or molecules across the vessel wall or an LEC monolayer, but they cannot discern whether molecules travel through junctions or via transcytosis, the movement of molecules through a cell. TEER studies use electrical resistance as an indicator of barrier integrity, but a 2020 study was unable to correlate to junctional changes with TEER results in a 2D brain endothelial cell model[13]. Furthermore, both assays can be influenced by gaps in the endothelial monolayer due to cell detachment rather than junction changes alone. . Collectively, these and other functional assays provide valuable measures of lymphatic transport and barrier integrity. However, since they only assess outcomes downstream of junction remodeling, they cannot directly link functional outcomes to morphological changes in endothelial junctions.

For lymphatic vasculature, evaluating total expression of junction proteins at the population-level has been used to show that junctions have formed[14,15] . Changes in permeability or barrier integrity observed in functional assays may be attributed to higher or lower junction protein expression, but total protein provides no context for junction morphology or spatial organization within or between cells[15,16]. Vascular endothelial (VE)-cadherin can be localized at the edge of cells to form long zipper-like junctions, resulting in inhibited flow. Alternatively, VE-cadherin can be localized internally via endocytosis during inflammation and eventually degraded or recycled by the cell[17]. Since LEC junctions are described qualitatively as continuous or linear for zip-per-like junctions and discontinuous or irregular for button-like junctions[1,2,6,7,16,18–20], visual changes in how and where VE-cadherin is expressed, as well as the associated junction morphology, can be leveraged for a more robust assessment of junction presentation. The benefit of using junction morphology to understand vascular function is that it allows researchers to establish structure-function relationships for a variety of junctional states. However, to establish definitive relationships, it is necessary to complement indirect measures and qualitative assessments with robust quantitative analyses that remove subjectivity and bias.

Over the years, efforts have been directed towards more quantitative methods for analyzing junction formation for lymphatic capillaries. In early lymphatics literature, methods for quantifying junctions included calculating the average number of junctions per unit area inside a lymphatic vessel, distance between junctions, and junction length[21,22]. These metrics leverage junction discontinuity as a key feature of button-like junctions by detecting discrete signals along a length of cell border. In addition, junctions are only measured with respect to button-like characteristics with no definitive profiles for zipper-like junctions. However, this framing may be attributed to button-like descriptions being reserved for lymphatic vasculature compared to blood vasculature. In recent years, the most used method of quantitatively characterizing endothelial junctions have been through thickness measurements, albeit the approach has not been exclusive to lymphatics. The Junction Analysis Program (JAnaP) quantifies junction organization at the borders of individual brain microvascular endothelial cells using Zonula Occludens (ZO)-1 signal to determine the ratio of junction length to junction thickness[23]. Junctions were classified as zipper-like and continuous (>15 pixel path length) or button-like, being either perpendicular (ratio >1.2) or punctate (ratio <1.2). We and other researchers have used thickness or width measurements to evaluate changes in endothelial junctions under varied substrate stiffnesses using CellProfiler/Fiji [24] and custom ImageJ/MATLAB pipelines[25,26]. Our study appeared to show thicker, more zipper-like VE-cadherin junctions with continuity in human dermal LECs on stiffer substrates, but the results were not combined with any functional tests[24]. Conversely, studies focused on blood vasculature observed wider junctions with increased vascular permeability (bovine and murine aortic endothelial cells) and leukocyte transendothelial migration (human umbilical vein endothelial cells) as stiffness increased [25,26]. Although those junctions were not classified in terms of button- or zipper-like morphologies, increased transendothelial movement of molecules or cells combined with more “jagged” presenting junctions suggest a more button-like morphology for cells on stiffer substrates. Despite progress in quantifying endothelial junctions, studies are still limited and are often applied to blood endothelial systems that do not distinguish the specialized button-like and zipper-like morphologies of LEC junctions. Existing lymphatic studies have provided a foundation for quantifying LEC junction morphologies based on thickness, distance, and border fragmentation, but focusing on singular features in isolation and continued reliance on qualitative classification does not capture structural changes across multiple metrics that better define junctional state. Therefore, a comprehensive and reproducible multi-metric approach is needed to quantify lymphatic junction morphology.

To advance approaches for quantifying LEC junction morphology, we developed a semi-automated image analysis pipeline that accounts for spatial expression of VE-cadherin junctions and visualization to generate distinct profiles that distinguish button-like junctions from zipper-like junction in human dermal LEC monolayers. The 3-step pipeline uses MATLAB (MathWorks, Natick, MA), CellProfiler (Broad Institute, Cambridge, MA), and Fiji (ImageJ) to process and analyze images of LEC-lined type I collagen channels within a microfluidic chip (Figure 1, Table A1, Table A2). Tumor necrosis factor (TNF)-α was used to disrupt LEC junctions toward a button-like morphology and represent a pathologically relevant cytokine. The pipeline accurately identified cell boundaries and used VE-cadherin signal and expression patterns to extract image-derived metrics that described VE-cadherin localization, object number, object size, branchpoints, endpoints, and thickness (Figure 1). To demonstrate its utility, we applied the pipeline to TNF-α treated and untreated LECs to detect changes in junctional morphology. Individual metrics identified distinct features of button-like and zipper-like junction morphologies, while Principal Component Analysis (PCA) integrated those measures into multivariate morphological profiles for each junctional state that expanded our interpretation beyond single-metric changes. Together, these findings demonstrate that the pipeline provides a standardized and adaptable approach for quantifying LEC junction morphology, thereby expanding the toolkit for evaluating in vitro lymphatic vascular models.

**Figure 1:**
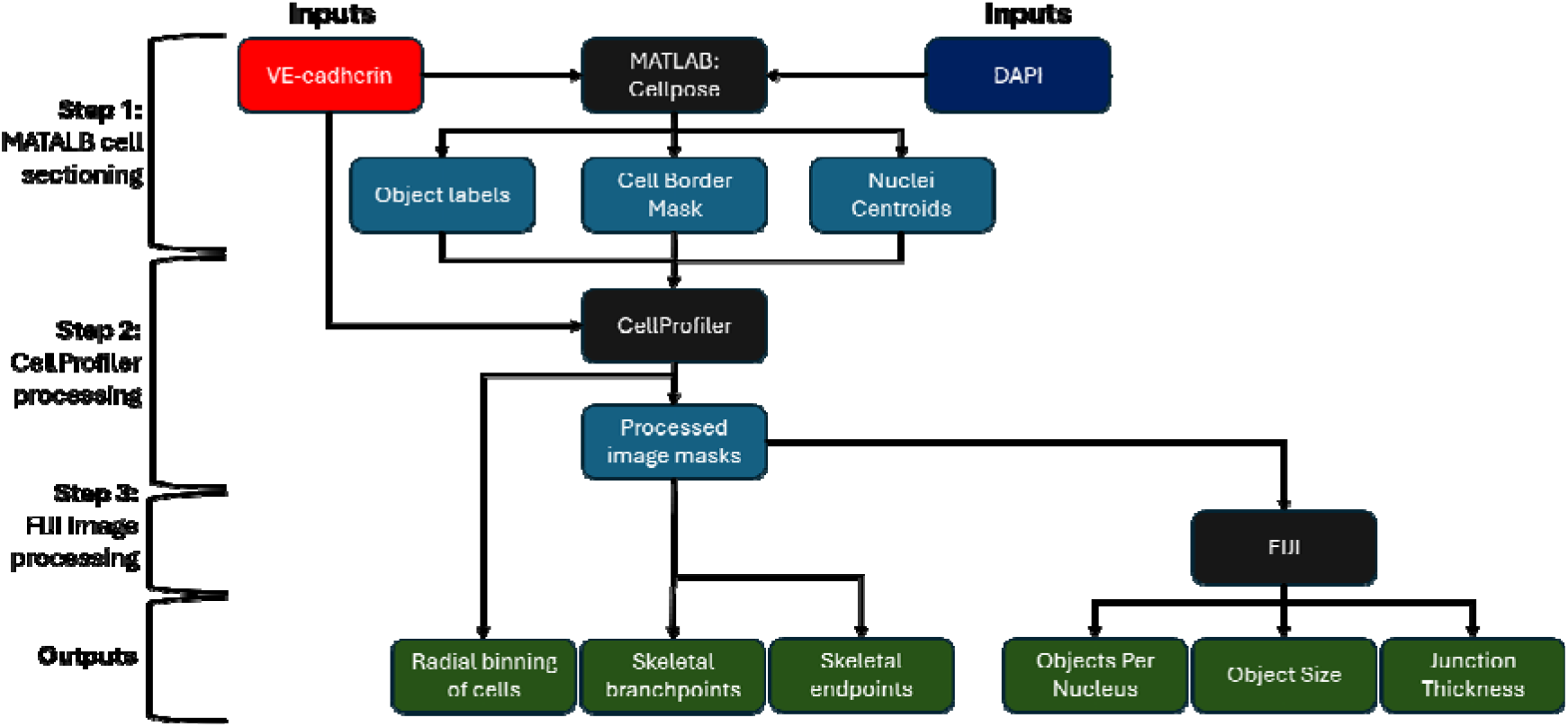
Image processing pipeline. The schematic shows the pipeline overview with inputs and outputs of each step of the process. Step 1) MATLAB was used to section the cells and identify cell borders from the original images (Cellpose). Step 2) CellProfiler™ is used to perform radial binning and image masking on original images and MATLAB outputs from Step 1. Image masks are directly used to identify and quantify skeletal branchpoints and endpoints. Step 3) FIJI is used to identify and quantify the number of objects per nucleus, object size, and junction thickness from image masks.

## 2. Materials and Methods

### 2.1 Lymphatic endothelial cell culture maintenance

Primary human dermal LECs (PromoCell, Heidelberg, Germany) were cultured and maintained according to the manufacturer using endothelial cell basal medium MV2 (MV2 media) (PromoCell) supplemented with fetal calf serum (5% v/v), epidermal growth factor (recombinant human, 5 ng/mL), basis fibroblast growth factor (recombinant human, 10 ng/mL), insulin-like growth factor (Long R3 IGF, recombinant human, 20 ng/mL), vascular endothelial growth factor 165 (recombinant human, 0.5 ng/mL), ascorbic acid (1 μg/mL), and hydrocortisone (0.2 μg/mL). Cells were maintained on tissue culture plastic with a 5 μg/cm^2^ fibronectin coating (Advanced BioMatrix, Carlsbad, CA, USA). For experimental samples, Modified MV2 medium was prepared similarly but without vitamin C and hydrocortisone additives to avoid anti-inflammatory and junction reinforcing effects[27–29]. Cells for this study were used between passages 6-11.

### 2.2 Microfluidic chip fabrication and sample preparation

Microfluidic chip devices with two parallel channels and a central extracellular matrix (ECM) chamber were fabricated according to Wang et al. [30] using a polydimethylsiloxane (Dow, Midland, MI, USA) (PDMS, 1:10 crosslinker:base ratio). The chips were plasma bonded to a coverslip then treated with Poly-L-lysine for 3 hours and a 0.05% v/v glutaraldehyde solution for 1 hour to promote ECM gel attachment before sterilization with ethylene oxide. Acupuncture needles (300 μm diameter) (Lhasa OMS, Weymouth, MA) were autoclaved and submerged in a 5% w/v bovine serum albumin solution (ThermoFisher Scientific) in 1X DPBS containing magnesium and calcium (DPBS++) (ThermoFisher Scientific) before being placed in devices. An acid-solubilized 8 mg/mL PhotoCol® (Advanced Biomatrix) a methacrylated type I collagen solution was neutralized(8% v/v neutralization solution) and combined with photoinitiator additives, 37.4 mg/mL Ruthenium (2% v/v) and 119 mg/mL Sodium Persulfate (2% v/v), yielding a final concentration of 7.122 mg/mL. Thirty microliters of neutralized PhotoCol®, was pipetted into the central ECM chamber and around the acupuncture needles. PhotoCol® gels with RuSPS and photo-crosslinking were leveraged to achieve pathologically relevant stiffnesses found in fibrotic microenvironments (∼6 kPa) [24]. Gels within the microfluidic chip devices were polymerized in a humidified incubator at 37°C with 5% CO_2_ for 60-90 minutes with the needles in place. After polymerization, the needles were removed to create two parallel channels in each device, and the gels were exposed to 405 nm light using an LED control system (ThorLabs, Newton, NJ, USA) for 90 seconds to generate stiff gels (Figure 2a). PhotoCol® gels were then washed with 1X DPBS++ 3-4 times over 24 hours on a rocker plate at room temperature protected from light to remove excess RuSPS.

**Figure 2:**
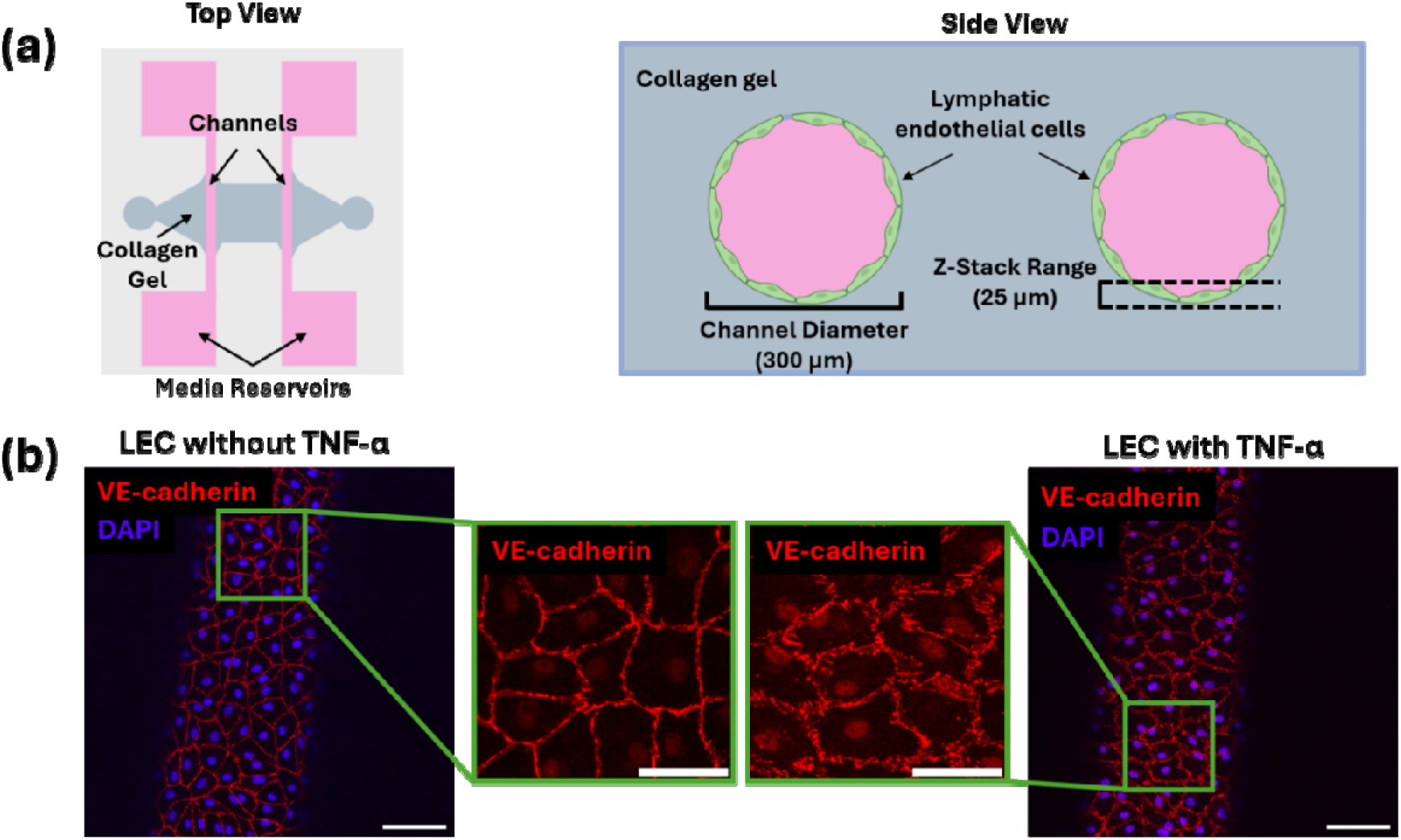
Visual representation of the 3D microfluidic device and in vitro model of a lymphatic vessel. (**a**) Schematic of the microfluidic device used to form a 3D lymphatic vessel in vitro. The top-down view of the device shows a 4 feature device with 2 channels per feature and a collagen gel reservoir. The side view shows the cross section of 300 µm diameter vessels in the device surrounded by a collagen gel. Dotted lines show the 25 µm z-stack thickness used for confocal imaging. (**b**) Representative images of z-stack maximum projections of LEC-lined channels without TNF-α treatment and with TNF-α treatment (red = VE-cadherin, blue = DAPI/nucleus). Green boxes show zoomed view of LECs to visualize junction features. Scale bar = 100 µm.

Cells were harvested from tissue culture flasks using a DetatchKit (PromoCell) and resuspend at 5×10^6^ cells/mL in Modified MV2 media. Collagen channels were seeded with 5 μL of cell suspension and inverted for 45 minutes in a humified incubator (37°C, 5% CO_2_) before repeating the seeding and inversion steps. After seeding, the media reservoirs (Figure 2a) were filled with Modified MV2 media before being placed on a rocker in a humidified incubator (37°C 5%, CO_2_). Media was changed daily for three days prior to adding media supplemented with 20 ng/mL TNF-α for an additional 24 hours to promote button-like junction formation. After four days of total culture time, samples were fixed and stained to visualize VE-cadherin and nuclei.

### 2.3 Immunostaining and image acquisition of VE-cadherin

For immunostaining, all incubation and wash steps were performed on a rocker plate, and multichannel pipette was used to avoid excess sheer in the vessels. Media was replaced with warmed pre-stain solution containing 1% (w/v) paraformaldehyde (ThermoFisher Scientific) with 0.01% (v/v) Triton-X100 (Sigma Alrich, St. Louis, MO, USA) in 1X DPBS++ and incubated at 37°C for 90 seconds. Pre-stain solution was replaced with warmed 4% (w/v) paraformaldehyde in DPBS++ to fix samples for 15 minutes at 37°C. Samples were washed with three gentle rinses of prewarmed DPBS++ at room temperature by pipetting up and down slowly. Warmed 0.5% (v/v) TritonX-100 was then added to each channel and incubated at 37°C for 10 minutes before three additional gentle washes. Samples were blocked with 2% (v/v) BSA in DPBS++ for 1 hour at room temperature or overnight at 4°C. Rabbit anti-human VE-cadherin primary antibody (ThermoFisher Scientific) was then prepared in the blocking solution (1:500), added to samples, and incubated at room temperature for 1 hour. Samples were washed with DPBS++ for 5 minutes per wash at room temperature, after which donkey anti-rabbit AlexaFluor 647 (ThermoFisher Scientific) secondary antibody prepared in blocking solution (1:1000) was added to samples and incubated for 45 minutes at room temperature. Samples were again washed with DPBS++ for 5 minutes per wash at room temperature before adding 4′,6-diamidino-2-phenylindole (DAPI) in DPBS++ (300 nM) as the nuclear counterstain for 10 minutes at room temperature. Three 5-minute wash steps using DPBS++ were used to remove excess DAPI prior to imaging. Imaging was performed using the 10X and 20X air objectives of a Leica Stellaris 8 confocal laser scanning microscope (Leica Microsystems, Nussloch, Germany). All images were acquired as 3D z-stacks at 25 μm total thickness (1 μm step size) to capture an appropriate image volume to generate maximum projection images. Fiji was used to scale images appropriately to ensure uniformity between 10X and 20X images, intensities for VE-cadherin images were rescaled as well for uniformity.

### 2.4 High throughput cell sectioning using Cellpose deep learning model

Before images could be analyzed, cell borders had to be identified and sectioned appropriately to ensure accurate results. To create a semi-automated and accessible model with higher throughput capabilities, we used a Python and MATLAB deep learning model called Cellpose that was trained on its internal cyto2 data[31,32] that included cell data and additional user contributed images[33]. Cellpose identified cell boarders using VE-cadherin images to create an object matrix, and MATLAB was used to find the border of the objects to create an object mask. A mask of the nuclei was then constructed through MATLAB using the DAPI (4′,6-diamidino-2-phenylindole) images, and exclusion criteria wereimplemented to remove errors in cell segmentation. Exclusion criteria began by removing any nuclei that were less than 20 pixels in area. Nuclei that spanned across cell boundaries were split, and cells containing more than one nucleus were not used in the pipeline. Finally, any cells that covered more than 15,000-pixel area were excluded to prevent open cells from being counted. Centroids of the accepted nuclei were identified and dilated radially by 5 pixels. Three outputs were generated from MATLAB: binary cell border, dilated nuclei centroid masks, and object labels (Figure 1). To validate cell sectioning the Dice Coefficient was used to compare overlap by applying an auto-threshold to VE-cadherin images to create VE-cadherin masks which were then overlayed onto cell boundary masks generated by Cellpose[34] . MATLAB’s Copilot was used to help synthesize the code to identify nuclei, implement exclusion criteria, and implement the Dice Coefficient.

### 2.5 Quantifying intensity distribution of internalized junction proteins using

CellProfiler Understanding spatial expression of VE-cadherin in cells provides important context for cellular response[10,35,36]. For VE-cadherin quantification, nuclei centroids were used to identify the cell nuclei within the “IdentifyPrimaryObjects” module of CellProfiler. Cell boundaries were defined from object labels generated by Cellpose using CellProfiler’s “ConvertImagetoObject” module. Prior to measuring VE-cadherin signal intensity, the intensities of original VE-cadherin images and VE-cadherin boundary masks from Cellpose were rescaled to cover the entire range of intensity (0 – 1; 0 = black, 1= white). Junction protein localization across the cell body was measured using the “MeasureIntensityDistribution” module to section cells into 10 bins radiating from and including the nucleus (Bin 1) to the membrane/cell edge (Bin 10). Higher intensity corresponds to lighter colors while lower intensity corresponds to darker colors.

### 2.6 Quantifying branchpoints, and endpoints using CellProfiler

Branchpoints and endpoints are metrics that can be used to describe junction shape and continuity. For accurate measurements for branchpoints and endpoints, images must first be skeletonized to reduce the width of foreground pixel shapes to a single pixel. Branchpoints are the pixels where dendrites, features exhibiting a branched or fractal pattern, split from the main body, creating pixels with more than 2 neighbors and a jagged shape that can indicate irregular cell boundaries. A skeletonized continuous zipper-like membrane would be visualized as a single pixel line outlining the perimeter of a cell. Endpoints are pixels with only a single neighbor and appear at the end of dendrites. Alternatively, they can be defined by gaps in cell perimeter that show discontinuity in the junctions, a clear sign of button-like junctions.

Cell boundary masks were dilated using disk size 5 and inverted. The “Smooth” function with the “Smoothed Keeping Edges” setting (edge intensity difference of 0.25) was then used to reduce speckling and ensure majority boundary signal was retained for further processing. In the next step, the “Morph” function was used to enhance lines of pixel length 5 or more and dampen internal cell structures in the image. By using a low pixel count of 5, continuous lines of both continuous junctions and fragmented button junctions could be enhanced. Next, the “Threshold” function was used with the following parameters: “Global”, “Minimum Cross-Entropy”, and threshold correction factor of 1 with a minimum pixel intensity of 0.05 images. Lastly, to ensure that the pipeline only measured membrane signal, the “ImageMath” function was used to subtract the inverted and dilated boundary image to remove the internal cell structures. The “MorphologicalSkeleton” function was used to reduce foreground membrane pixels to lines of a single pixel width which is necessary for the next step in the pipeline. The “MeasureImageSkeleton” function was then used to measure branchpoints and end-points. For final quantification, branchpoint and endpoint measurements were normalized to the number of identified centroids.

### 2.7 Quantifying objects per cell and object size using Fiji

Object count and size metrics are direct indicators of junction discontinuity and fragmentation. When junctions form continuously around a cell, a large single object wraps around the perimeter of a cell. When junctions fragment, they are visualized as small individual separate points of connection between cells. Using “Analyze Particles” in Fiji, the number of membrane objects can be counted with automatic calculation of size averages per image. The images used in Fiji were the binary images that originated from CellProfiler and were created before skeletonizing to measure branchpoints and end-points. To determine the number of objects per cell, the “Analyze Particles” tool (minimum pixel count = 5; maximum pixel count = Infinity) was applied to the images. The pixel length used for the “Morph” function in CellProfiler was used to determine the minimum size threshold for the “Analyze Particles” tool in Fiji.

### 2.8 Quantifying junction thickness using Fiji

Junction thickness is a common metric that is used to describe endothelial junctions and is often correlated with barrier integrity for endothelial cell populations[24,37]. Fiji’s Vessel Analysis plugin was used to measure junction thickness[38]. The plugin works well for continuous junctions, because they are similar in shape to vascular networks. For discontinuous junctions, continuity was created using a sequence of Fiji functions that were applied to all images: 1) Close, 2) Dilate, 3) Close, 4) Erode. The geometry to distance map tool (Fiji) was used to perform a distance transformation on the foreground objects and overlay a heatmap onto the mask.[38]. Once the distance map was created, the Vessel Analysis plugin (Fiji) was used, and the “Diameter Measurements” tool converted the distance map into an average thickness measurement across the entire network of junctions.

### 2.9 Data Analysis

Data was collected from two independent experiments. The non-treated groups included N = 12 (number of image samples) and n = 36-66 (number of cells) across all trials, and TNF-α treated groups contained N = 6 and n = 32-59 across all trials. Image sample averages were calculated and input into GraphPad Prism 11 (GraphPad Software Inc., La Jolla, CA, USA) for univariate and multivariate statistical analyses. For each metric, normality of the data was tested using a Shapiro-Wilk test (α = 0.05). For spatial localization of VE-cadherin signal, a two-way ANOVA showed significant main effects (treatment and bins) and significant interaction. Therefore, a post-hoc Sidak’s multiple comparison test was used to evaluate simple effects within each treatment group and within each bin. A Welch’s t-test was used to compare independent sample means between TNF-α treated and non-treated groups for branchpoints, endpoints, objects per cell, object size, and junction thickness. Results were considered statistically significant at * p < 0.05, ** p <0.01, **** p<0.0001. Principal component analysis (PCA) was performed on the morphological metrics to evaluate multivariate relationships between TNF-α treated and non-treated groups. Principle components were retained until approximately 90% of the total variance was explained, and PCA scores and biplots were used to distinguish groups and identify contributions from variables.

## 3. Results

### 3.1. Cellpose deep learning model produces reliable cell sectioning for lymphatic endothelial cells (LEC)

Representative outputs (Figure 3a) of Cellpose generated cell boundaries and MATLAB identified nuclei were validated visually by merging the DAPI and VE-cadherin input images (Figure 3b, RGB Composite). Correct boundary and nuclei identification were determined qualitatively by observing the overlays of VE-cadherin signal with the boundary mask and the DAPI signal with the centroids (Figure 3b, bottom image). Segmentation was verified using the Dice Coefficient which showed a slight between treatment groups, but differences were not significant (Figure 3c). When verifying segmentation, values between 0.7 -0.9 are classified as “good” segmentation with significant overlap whereas values below 0.6 are considered to be poorly segmented[39,40] Cells without TNF-α, representing zipper-like morphologies, scored higher on average (0.81) than TNF-α treated cells that were more button-like (0.77). The slightly higher average Dice Coefficient for zipper-like junctions (untreated cells) was due to increased junction continuity in the mask of the VE-cadherin signal. Since both Dice Coefficient values were close to 0.8, segmentation was deemed acceptable. However, since visual differences between the thickness of the VE-cadherin junctions and the Cellpose generated boundary mask were observed, a dilation step was applied to the Cellpose mask to ensure adequate overlap.

**Figure 3:**
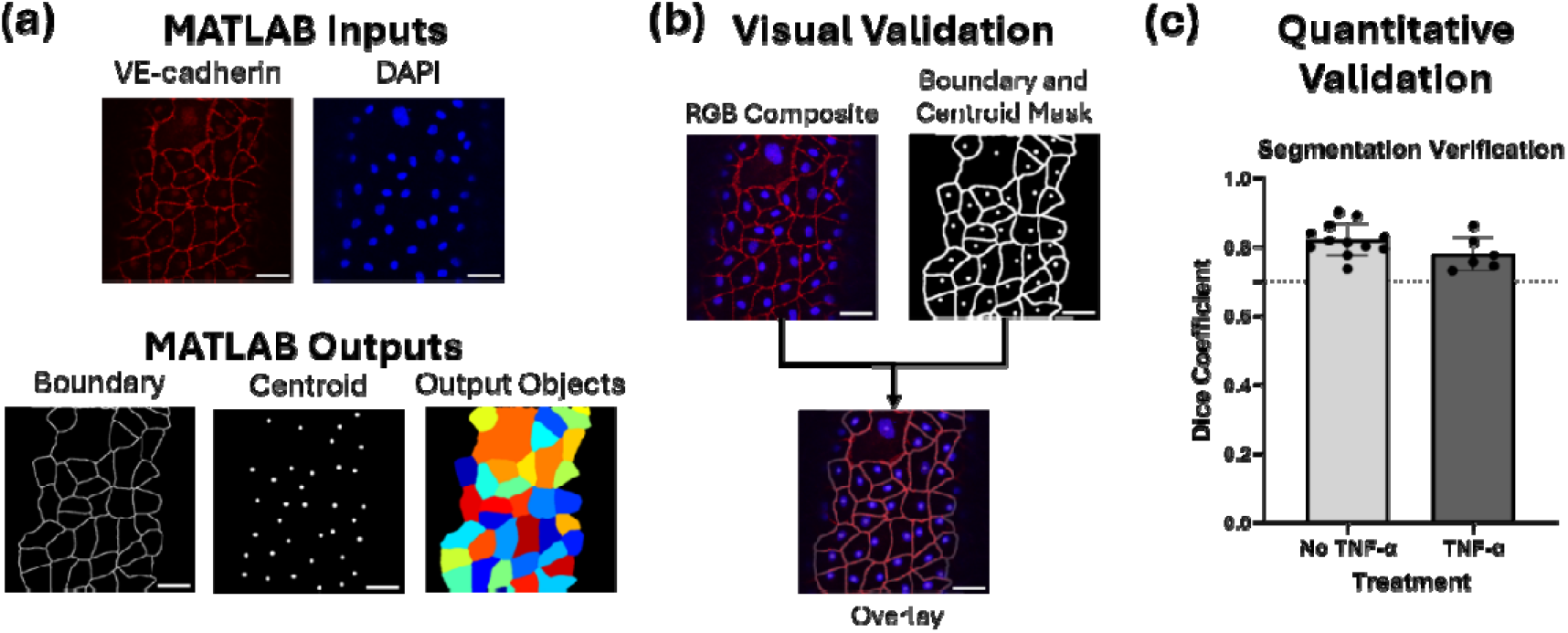
Inputs, outputs, and validation for Cellpose deep learning model. (**a**) Input images and output masks generated via Cellpose (MATLAB) to create cell boundaries (VE-cadherin signal, red) and centroids (DAPI/nuclear signal, blue) for further processing in CellProfiler. (**b**) Qualitative comparison of the merged input image channels (top left) and nuclei centroid mask and cell boundary mask using FIJI (top right). The masks are then overlayed on the merged image channels (bottom) to show overlap between the MATLAB generated boundaries and nuclei and the true cell boundaries and nuclei Scale bar = 50 µm. **(c)** Dice coefficient representing the quantitative assessment of the degree of overlap between the original images and generated masks used for analyzing LEC without TNF-α (light gray) and with TNF-α (dark gray) Data are represented by scatter bar graphs with mean±SD.

### 3.2 Radial intensity measurements detect differences in VE-cadherin spatial localization between button-like and zipper-like morphologies

VE-cadherin signal intensity within in LECs was measured radially from the nuclei to the membrane edge across 10 bins to determine areas of high and low intensity (Figure 4a). Measurements from bins 1-3, covering the nuclei and area immediately surrounding the nuclei, showed a slight increase in mean intensity in groups containing TNF-α compared to groups without TNF-α with a significant increase in bins 2 (* p < 0.05) and 3 (** p < 0.01) (Figure 4b). Bins 4-9 saw a decrease in overall intensity compared to the bins at the nucleus and cell membrane with no statistical difference between treatment groups, but the intensity of TNF-α treated cells were slightly higher. As expected, signal intensity decreased significantly with TNF-α treatment in bin 10 (**** p<0.0001) showing that less VE-cadherin localized to the cell border in those cells (Figure 4b). Given that TNF-α treatment induces button-like junctions, these results show localization away from the cell membrane and toward the nucleus and surrounding cytoplasm when junctions follow a button-like morphology. When cells exhibited a zipper-like morphology without treatment, the VE-cadherin signal increased accordingly at the cell membrane. The observed trends in VE-cadherin localization associated with zipper and button-like junctions demonstrate that radial intensity of VE-cadherin signal can be used to track protein localization during junction remodeling and transitions to gain morphological insights.

**Figure 4:**
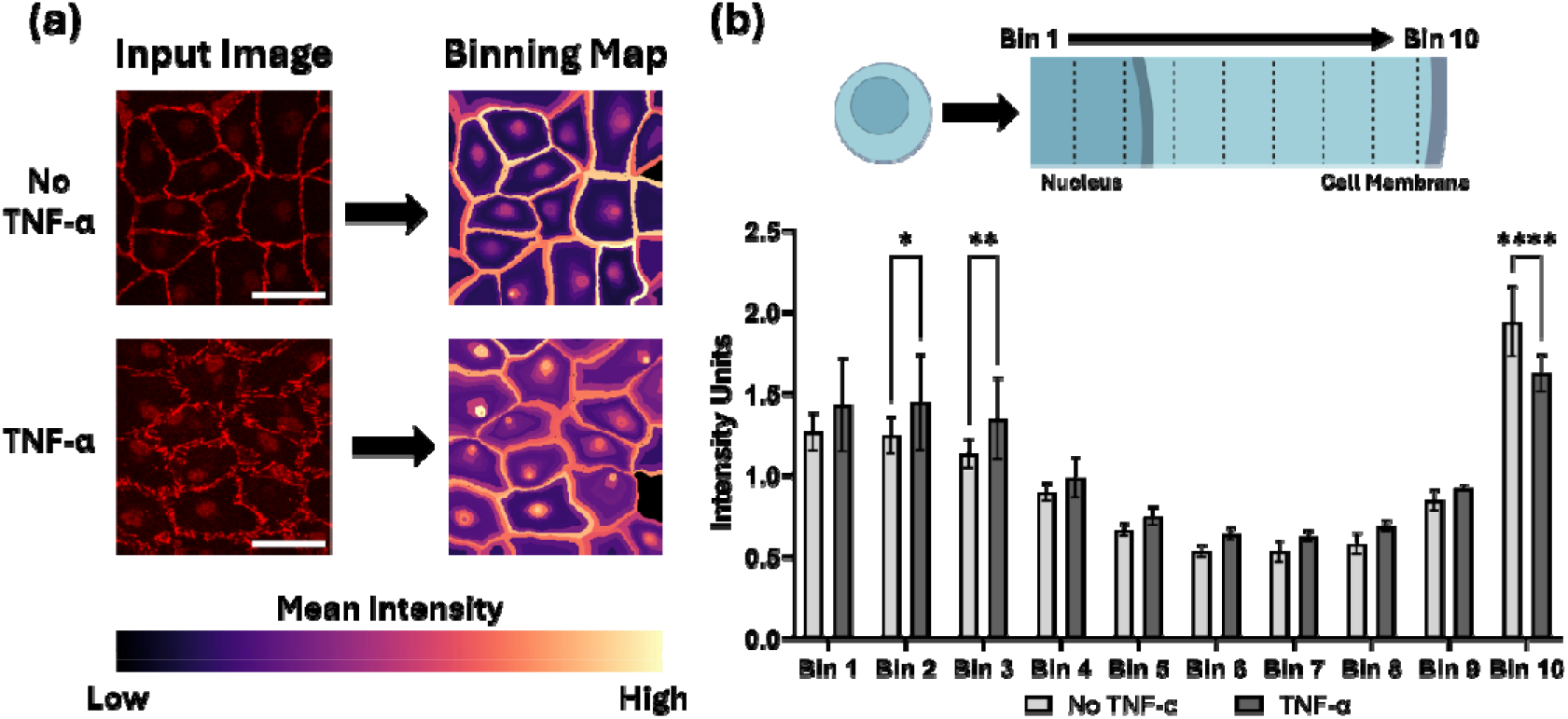
Quantification of VE-cadherin localization in lymphatic endothelial cells. (**a**) Illustrative example of VE-cadherin image input and binning output shown as a distance heatmap of low to high mean intensity (Scale bar = 50 µm). (**b**) Schematic of binning orientation for Bins 1-10 radiating from the cell nucleus to the cell border and corresponding quantification of VE-cadherin intensity across the cell bodies of LEC without TNF-α (light gray) and with TNF-α (dark gray). Bars represent mean±SD. Metrics were compared with 2-way ANOVA using Šídák multiple comparisons for independent samples (* p < 0.05, ** p <0.01, **** p<0.0001). Bin schematic generated using Biorender™.

### 3.3 Branchpoint and endpoint quantification capture differences in junction continuity between button-like and zipper-like morphologies

To measure the number of branchpoints and endpoints in the junctions, images were processed to visualize junctions and skeletonize to shrink image shapes down to a single pixel width feature (Figure 5). A zoomed image of skeletonized VE-cadherin (Figure 5a) in LEC without TNF-α treatment shows junctions following a linear path with few branching points and fewer endpoints. When junctions become more button-like with TNF-α treatment, dendrites form along the linear path creating an irregular shape with numerous endpoints. There was a trend for increased branchpoints and endpoints when junctions were disrupted with TNF-α treatment. Branchpoints per cell increased significantly from 35.6 ± 17.1 to 88.9 ± 31.7 branchpoints (** p < 0.01) while endpoints increased significantly from 20.7 ± 8.8 to 50.0 ± 7.2 endpoints (**** p< 0.0001). Branchpoints and endpoints both positively correlated with button-like junction morphology and are inversely correlated to zipper-like junction morphology, demonstrating their ability to distinguish junction morphologies individually and jointly.

**Figure 5:**
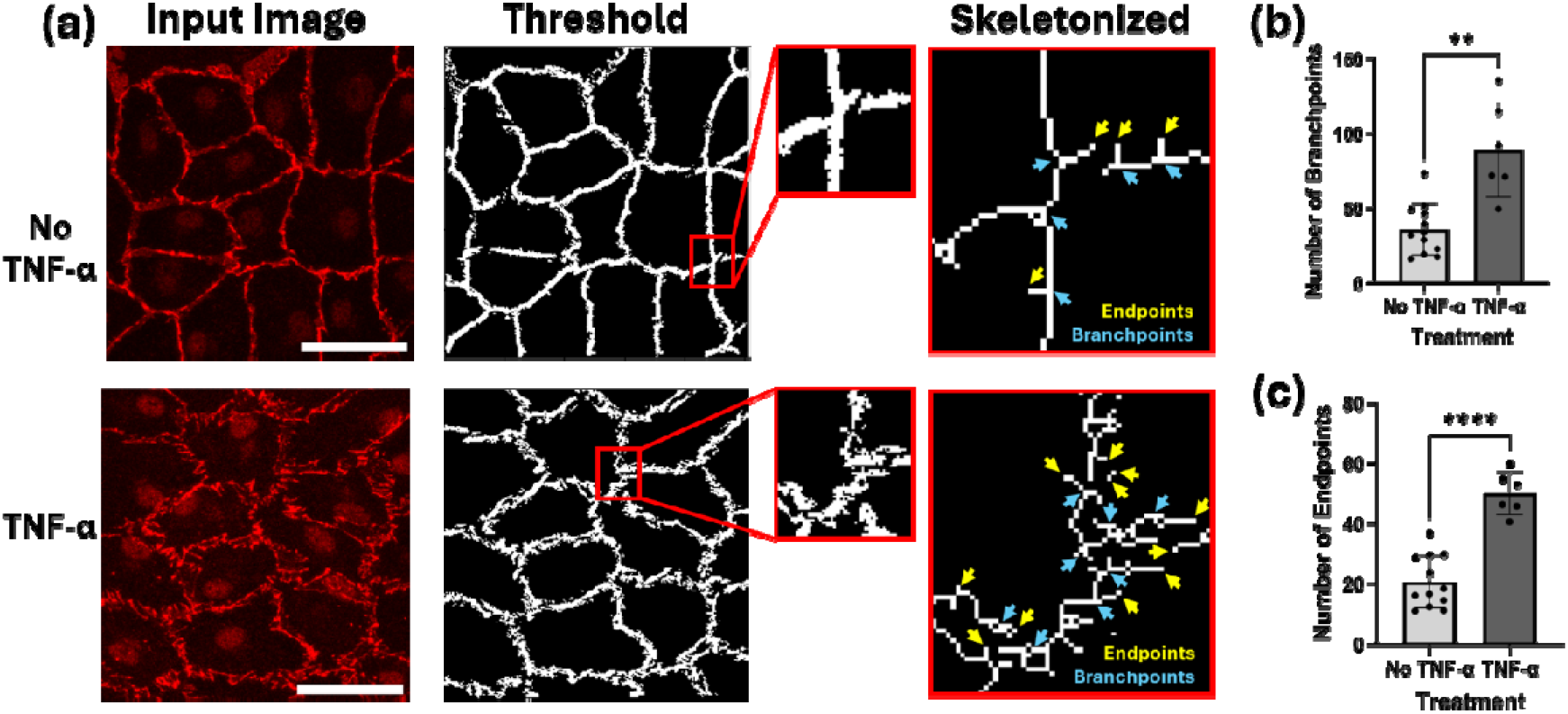
Quantification of branchpoints and endpoints. (**a**) Visual representation of image processing from image input to thresholded CellProfiler mask and skeletonization (Scale bar = 50 µm). Red boxes represent the zoomed portion of the threshold image with corresponding branchpoint and endpoints in the skeletonized image. (**b**) Number of branchpoints and (**c**) Number of endpoints (normalized by the number of cells per image) for LEC without TNF-α (light gray) and with TNF-α (dark gray). Data are represented by scatter bar graphs with mean±SD. Metrics were compared with Welch’s t-test (** p <0.001, **** p<0.0001).

### 3.4 An inverse relationship between objects per cell and object size describe differences in junction fragmentation between button-like and zipper-like morphologies

To quantify objects per cell and object size, the “Analyze Particle” function in Fiji was used on masks generated from CellProfiler (Figure 6a). There was an increasing trend in VE-cadherin disruptions or fragmentation indicated by an increase in the number of objects per cell in the presence of TNF-α treatment (Figure 6b). In groups without TNF-α treatment there was an average of 1.03 ± 0.08 objects per cell while groups with TNF-α treatment were significantly higher at an average of 9.4 ± 0.7 objects per cell (Figure 6b) (** p < 0.01). For the object size, an inverse relationship was observed with significantly larger objects for cells without TNF-α treatment at 353.7 ± 169.2pixel area compared to 152.6 ± 85.11 pixel area for cells with TNF-α treatment (Figure 6c) (** p < 0.01). Results show that the pipeline can quantify objects per cell and object size and both metrics be used to distinguish TNF-α associated buttoning of junctions from more zipper-like junctions.

**Figure 6:**
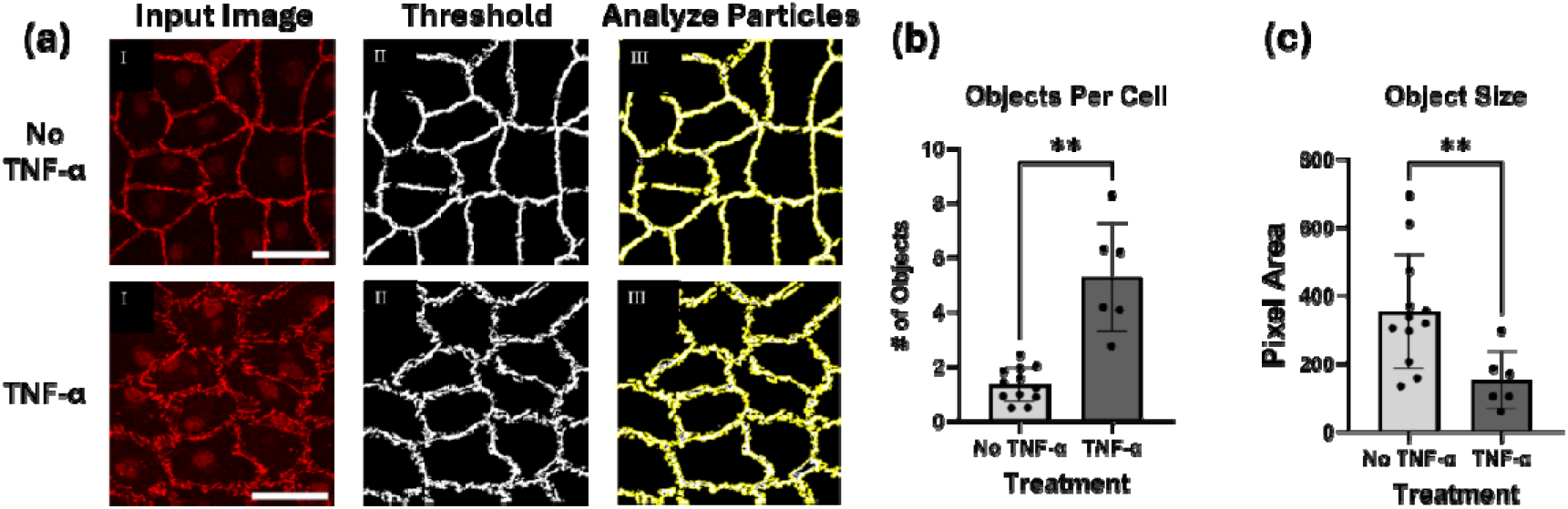
Quantification of objects per cell and object size metrics. (**a**) Illustrative example of process from original VE-cadherin image (left) to thresholded CellProfiler mask (middle) and final particle analysis using FIJI (right) (Scale bar = 50 µm). (**b**) Counted number of objects (normalized by the number of cells per image) for LEC without TNF-α (light gray) and with TNF-α (dark gray). (**c**) Average pixel area representing object size for treated and non-treated groups. Data are represented by scatter bar graphs with mean±SD. Metrics were compared with Welch’s t-test (** p <0.01).

### 3.5 Vessel Plugin Detects Morphology-Associated Differences in Lymphatic Endothelial Junction Thickness

Junction thickness was measured using the Vessel Analysis plugin (Fiji) applied to image masks generated from CellProfiler. The initial processed images appear to show thicker lines for cells with TNF-α treatment (Figure 7a). When quantifying thickness, LEC with TNF-α treatment were significantly thicker at 1.22 ± 0.13 μm versus 1.03 ± 0.78 μm for LEC without TNF-α treatment (Figure 7b) (* p < 0.05). The analysis plug-in shows the ability to differentiate between treatment groups and related junction buttoning or zippering based on junction thickness.

**Figure 7:**
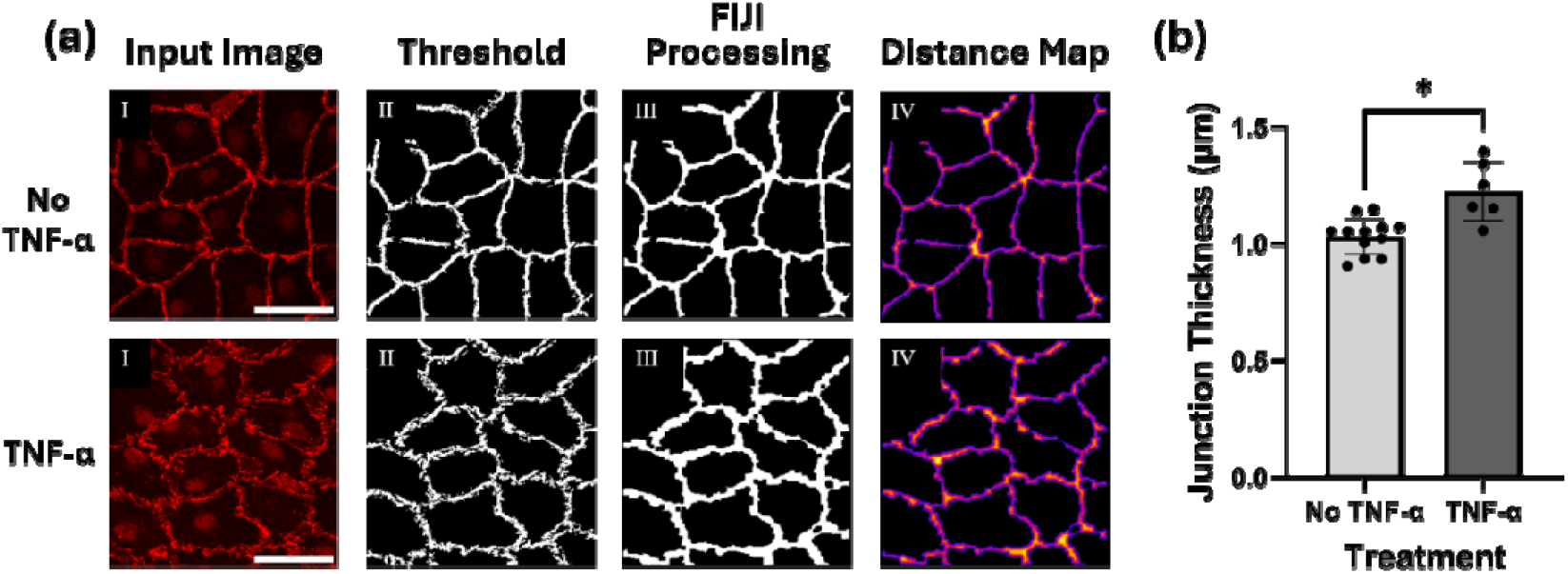
Quantification of junction thickness metrics. (**a**) Illustrative example of process from original input VE-cadherin image (I) to thresholded CellProfiler mask (II), FIJI Processed mask (III) and final Distance Map (IV) for quantification (Scale bar = 50 µm). (**b**) Junction thickness of VE-cadherin junctions in LEC without TNF-α treatment (light gray) and with TNF-α treatment (dark gray). Bars represent mean±SD of junction thickness calculated with the “Vessel Analysis” plugin in FIJI. Data are represented by scatter bar graphs with mean±SD. Metrics were compared with Welch’s t-test (* p <0.05).

### 3.6 Principal component analysis (PCA) identifies distinct VE-cadherin morphological signatures

Principal component analysis (PCA) was used to evaluate relationships among all metrics in this study and determine how the individual metrics combine to define distinct morphological states. The first two principal components captured 90.7% of the total variance in the dataset, with PC1 accounting for 64.8% (eigenvalue = 3.89) and PC2 accounting for 25.8% (eigenvalue = 1.55). The sample scores showed distinct separation between TNF-α treated and untreated onto PC1 (Figure 8). TNF-α treated samples clustered along the negative direction of PC1, and untreated samples were prominently distributed along the positive direction. The negative loading for TNF-α treated samples is driven by higher values for object number, endpoints, branchpoints, and thickness, indicative of button-like junctions for individual metrics. When these metrics were considered jointly, there is a potential signature for button-like morphology. The samples without TNF-α treatment aligned with increases in object size and VE-cadherin localization at the cell membrane which differs from button-like trends and is more characteristic of zipper-like junction morphology. Variation along PC2 further separates the samples on object number. Together, these multivariate results demonstrate that TNF-α treatment produced a shift from a more continuous signal at the LEC barrier that was associated with zipper-like junctions, to disrupted and fragmented junctions with more button-like characteristics.

**Figure 8:**
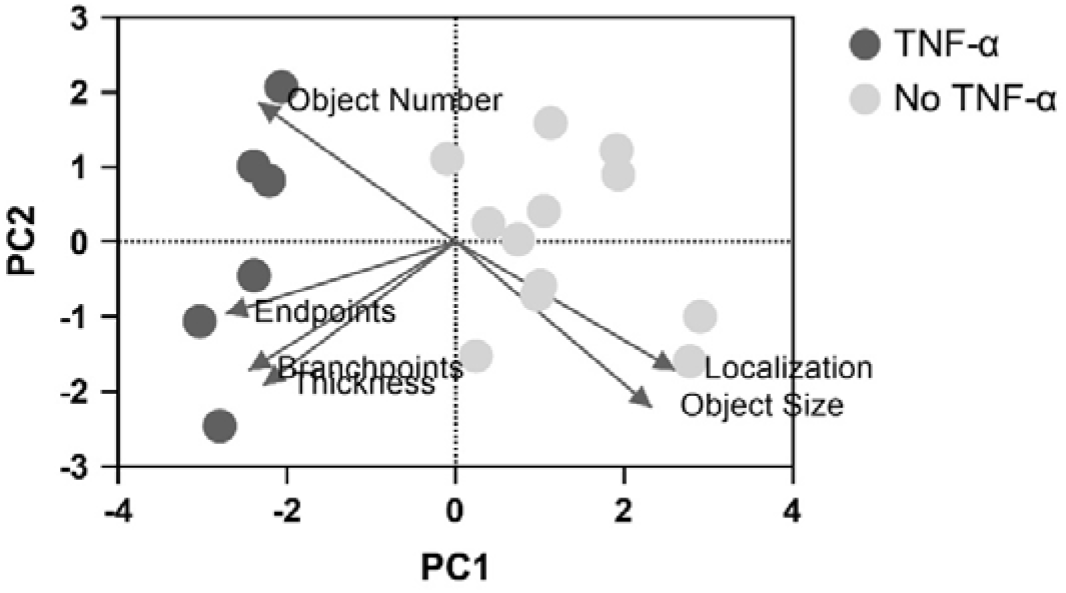
Principal component analysis biplot of junction metrics. Biplot showing the distribution of untreated control (No TNF-α, light gray) and TNFα treated (dark gray) LECs. Points are positioned according to their scores on principal component 1 (PC1) and principal component 2 (PC2) with overlaid loading vectors (lines with arrow) for quantified junction metrics: VE-cadherin Localization, Object Size, Object Number, Endpoints, Branchpoints, and Thickness. Arrow direction indicates the direction of increasing values for each metric in PCA space. Arrow length reflects the relative contribution to the components.

## 4. Discussion

In this study, we developed a 3-step image analysis pipeline with multiple metrics, new and existing, to quantify differences between VE-cadherin button-like and zipper-like junction morphologies in small diameter lymphatic vessels. This analysis tool was implemented on an in vitro lymphatic-on-a-chip platform prepared with human dermal LECs that were cultured within a stiff collagen gel channel with and without TNF-α treatment to generate zipper-like and button-like junction morphologies, respectively. Our work sought to expand lymphatic junction analysis beyond basic qualitative assessment and solo metric analysis to capture comprehensive junction information about lymphatic junctions. In addition to existing metrics such as junction thickness and object size, we identified and quantified new metrics that included VE-cadherin localization, objects per cell, branchpoints, and endpoints. These metrices can be evaluated individually or in combination to describe LEC junctions and classify them as button-like or zipper-like based on a morphological signature. Results of the study show that our new analysis approach quantifies metrics that describe both button and zipper-like junctions that reflect varied pathophysiological conditions. Collectively, these findings support the use of this new analysis pipeline as a useful tool to study junction morphology in the lymphatic endothelium in response to junction altering stimuli (e.g. inflammation).

Prior to quantifying junction metrics, we verified adequate cell sectioning, border identification and identified centroids of cell nuclei (Figure 3). Cellpose was for cell sectioning over the built-in toolbox in CellProfiler, because Cellpose excels at identifying cell boundaries across a range of imaging modalities with a comparatively low learning curve and an ability to section images consistently regardless of junction morphology[31]. However, centroid identification was performed outside of the Cellpose model to limit overlapping nuclei and cell boundaries issues that could impact intensity distribution steps for VE-cadherin localization analysis. Cellpose accurately identified cell boundaries within the LEC monolayer which was confirmed visually and with Dice Coefficient values of 0.77 and 0.81 that describe a “good” fit. This verification also showed that Cellpose could determine which cells would be counted towards analysis. However, the pipeline is adaptable enough that if incorrect boundaries or centroids were ultimately observed, users have an opportunity to customize the code to suit their model.

Each metric identified and quantified from our pipeline describes LEC junction morphology through a different perspective. While some metrics can classify junctions as button-like or zipper-like on their own, combining metrics strengthens the analysis and contributes more to the assessment. VE-cadherin localization using radial intensity measurement was achieved with binning across the cell body. Often, the focus of VE-cadherin expression is solely at the membrane, which misses critical information about VE-cadherin dynamics. Internalization of VE-cadherin protein causes the zipper-like junctions between cells to segment creating more button-like junctions [29,30]. With internalization, signal intensity is expected to shift away from the periphery of the cell membrane and increase towards the cytoplasm and nucleus, which creates an opportunity to leverage that shift as an indicator of junctional state. We observed the most significant increase or VE-cadherin signal closer to the nucleus (Bins 2 and 3) with reduced signal at the cell membrane in the presence of TNF-α, classifying this expression pattern as associated with button-like junctions (Figure 4). Conversely, VE-cadherin signal increased significantly at the cell membrane (Bin 10) without TNF-α, suggesting more zipper-like morphology. Collectively, this result demonstrated successful quantification of TNF-α induced changes in VE-cadherin localization, providing information about VE-cadherin signal beyond the cell membrane and corresponding to junctional state. Successful tracking of VE-cadherin localization across the cell body, rather than a binary assessment of cytoplasm versus cell membrane, was possible due to segmenting the cell radially outward from the nucleus. Choosing the appropriate number of bins was critical to the analysis. If too few bins were chosen, the signal at the cell membrane would be diminished by the cytoplasmic signal also captured within that bin. Too many bins would create many narrow regions with not enough pixels. Ten bins gave sufficient spatial resolution to distinguish signal intensity across the cell and detect clear trends of localization at the cell membrane and the perinuclear space. Overall, the binning method with radial intensity measurement shows that the pipeline goes beyond cell membrane expression to capture VE-cadherin dynamics related to localization and offer insights into changes in LEC junction morphology between button and zipper. In addition, while radial intensity measurements showing protein expression and localization can be valuable individually, their value increases when combined with additional metrics such as object size metrics.

Objects per cell and object size were both quantified to individually capture junction fragmentation and distinguish button-like junctions from zipper-like junctions. However, the two metrics are related. To avoid any discrepancies due to cell number per image, object per cell and object size counts were divided by the number of cells or nuclei in the image. We hypothesized that more junction fragmentation due to buttoning would produce more objects that were smaller in size. The overall result showed that TNF-α treatment significantly increased the number of objects per cell approximately 5-fold, which reflects more junction fragmentation and buttoning (Figure 6). As expected, object size decreased with TNF-α treatment, while samples without TNF-α treatment exhibited an opposite trend of fewer objects that were approximately 2 times larger than treated samples. The inverse trend seen in object count and size are clear indicators of the degree of junction fragmentation. When combined with VE-cadherin localization these metrics provide a quantitative measure of the cellular response by relating the intensity of internalized VE-cadherin around the nucleus to the degree of junction fragmentation at the cell boundary.

Beyond localization and object metrics, skeletal branchpoints and endpoints were additional metrics used to measure junction shape, fragmentation, and monolayer continuity to distinguish button-like and zipper-like junctions. Branchpoints are identified as a pixel with more than two neighbors in the foreground indicating an intersection of two or more lines. Endpoints are identified as pixels in the foreground with only one other neighboring foreground pixel. For branchpoint and endpoint measurements, normalizing the total counts to the number of cells identified in each image was crucial to overcome cell count variability in each image. The original hypothesis assumed that branchpoints would only be observed in regions where three or more LECs met, creating an intersection of junctions with continuity of the monolayer. A drop in the number of branchpoints would be seen as a decrease in intersections as junctions became button-like and cells separated. However, the results told a different story (Figure 5). Without TNF-α treatment, monolayer continuity was largely maintained with fewer endpoints, consistent with zipper-like morphology and few branchpoints at cell intersections. With TNF-α treatment, we expected to see a decrease in branchpoints and an increase of endpoints due to VE-cadherin fragmentation that were thought to represent junction discontinuity observed with a more button-like morphology. Although increased junction discontinuity was observed with TNF-α treatment, the jaggedness of the LEC membrane yielded more branchpoints along the junction boundaries with more end-points. Together, these membrane irregularities captured changes that align with button-like junction morphology in TNF-α treated samples. Moreover, the metrics themselves provide a way to describe junction continuity observed across the monolayer and for single cells (Branchpoints and Endpoints) and fragmentation at the cell membrane (Endpoints).

While junction thickness results showed significant differences between treatments, we do not believe the metric can uniquely describe button-like or zipper-like junctions. Studies performed by other research groups linked increased junction thickness to higher vascular permeability [23,37], suggesting that thicker junctions might be indicative of button-like morphologies based on function. However, these studies did not seek to explicitly classify junctional state. Our results had a similar outcome in that cells with TNF-α treatment and button-like characteristics consistently had greater junction thickness (Figure 7), but junction fragmentation likely played a role in our outcome due to image processing. The Vessel Analysis plug-in used to calculate thickness requires continuity to perform the measurement, so images were processed to artificially make the cell membrane more continuous. To ensure that all junctions were handled the same, all images were processed using the same functions regardless of initial degree of discontinuity. The endpoint visualization represented in Figure 5 shows the extent of fragmentation in TNF-α treated cells. As those endpoints were connected and smoothed in preparation for thickness measurements, the resulting image mask was unsurprisingly thicker. Given that our result established fragmentation as a feature of button-like junctions, one could argue that button-like junctions are actually “thicker”. However, the thickness is artificial and likely does not affect true junction structure. Therefore, even though our pipeline can detect junction thickness differences between button-like and zipper-like junctions in our study, the metric alone is not sufficient to classify junctional state.

When taking all metrics into account, PCA further demonstrates the value of classifying LEC junction morphology as a coordinated multivariate phenotype rather than individual metrics. The separation between TNF-α treated and untreated samples showed that metrics followed opposite patterns. Button-like junctions with TNF-α treatment were associated with increased object number, branchpoints, endpoints, and thickness, dominated by junction fragmentation and discontinuity. Conversely, zippered LECs without TNF-α treatment exhibited more junction continuity with larger objects and more VE-cadherin localization at the cell membrane. These distinct signatures highlight key differences between junction morphologies and reinforce the strength of combining complementary metrics. Because PCA uses quantitative outputs generated by the analysis pipeline, it also provides a strategy for comparing lymphatic junction morphologies across conditions and experimental models in future studies. The analysis also provides an opportunity to explore morphologies across the button-to-zipper spectrum to further investigate lymphatic junction dynamics.

As we place this work within the context of other approaches to quantify LEC junction morphology, some key observations were made. When compared to previous studies that measured junction thickness, junction morphology was either paired with a permeability assay or biochemical effect on junction thickness was not the focus of the study[24,37,42]. Junction thickness is limited in its ability to classify junction morphology on its own and is better suited to compare basic visual differences in junctions. On the other hand, junction count per vessel surface area, junction length, and distance between junctions were some of the first real methods of quantifying junction morphology and describing monolayer level differences in junction morphology [21,22]. These metrics are most closely related to the objects per cell and object size metrics described in the current study which showed similar trends in junction size and count metrics for LEC with button-like junctions. A few limitations of these methods was their lack of automation, translating junctions per vessel surface area to a 2D system, and assumptions for consistent cell shape and size when comparing changes in cell morphology. Dynamic changes in cell shape impact the junctions per area measurement, because vessel or 2D area alone does not fully take cell shape differences into account and depends on a continuous monolayer. To accurately use vessel or 2D monolayer area to count junctions and compare between groups, LEC must have the same cell area when oak leaf shaped or elongated. There must also be the same or similar number of cells between groups, and no gaps from detached cells must be present. Our method removes the assumption for equal cell area between treatment groups by normalizing to number of cells in an image. Beyond identifying more comprehensive metrics to describe LEC junction morphology, we sought to make morphological analysis more automated than existing approaches. The JAnaP pipeline excels at cell junction morphological analysis on individual cells and was initially validated in brain endothelial cells by measuring changes in ZO-1 junction shape and count per cell in response to varied stiffness and cAMP [23] . However, JAnaP is not designed for bulk monolayer characterization and is semi-automated, requiring manual sectioning of individual cells and small datasets with few cells. Given that the creators of the JAnaP pipeline acknowledged the limitations on quantitative analysis of protein presentation and subsequent assessment of healthy physiology and pathology, we see the increased automation of our pipeline as a way to bridge that gap by being more suitable for larger data sets and having potential to work in tandem with programs like JAnaP to comprehensively analyze LECs.

Beyond the pipeline itself, culture format was another important element to our study given the architecture of native lymphatic vessels. Morphological analysis was conducted using a 3D microfluidic chip [30] designed to recreate a cylindrical lymphatic vessel in a fibroinflammatory environment with a stiffened collagen matrix and TNF-α treatment. This format differs from many studies of LEC morphological analysis that culture LECs on a 2D planar surface [43,44]. The advantage of the chip model is that LEC can be exposed to curvature reminiscent of a lymphatic vessel while being entirely surrounded by a collagen gel with tunable biophysical properties. Previous studies have shown in different endothelial cell populations that vessel-like curvature can alter the morphology across endothelial subtypes[45–47]. For LECs specifically, tubular channels elicited lower circularity and greater elongation versus rectangular channels, when alignment was measured. Vessel shape also influenced LEC alignment along with flow[47]. Together, these findings suggest that the curved geometry of the lymphatic vessel may contribute to LEC morphology and organization in ways that are not captured by conventional 2D culture, providing an important consideration when comparing junctional phenotypes across culture formats. Moreover, while we acknowledge that the microfluidic lymphatic-on-a-chip model is not an exact replica of in vivo lymphatic vasculature, it serves as an appropriate in vitro platform to bridge the gaps between 2D in vitro cultures and 3D in vivo tissues.

Although our new analysis pipeline expands the ability to evaluate LEC junction morphology quantitatively, there are still some limitations. Even though the Dice Coefficient used to compare original images and image masks was in the “good” range, we believe that smoothing at the cell borders prevented a higher Dice Coefficient. Smoothing limitations were tolerated, because segmentation visually overlapped well enough with the VE-Cadherin signal to capture junction changes. In addition, the dilation function we added to the pipeline became standard procedure for the approach. An additional limitation was noted for junction thickness. Placing junction thickness measurements within the appropriate context was a challenge as previously described. The complexities of junction structure, particularly for button-like junctions, combined with heavy image processing required by the Vessel Analysis plug-in generated image masks that were artificially thick or thin. Although all images were processed the same, an alternative approach for measuring thickness should be considered. For multivariate analysis, PCA summarized variation among the chosen metrics but did not offer mechanistic insight into how junctions change between button- and zipper-like or provide information into thresholds for either morphology. Moreover, our analysis does not capture the spectrum of button-to-zipper junction and only classifies them into binary categories. Despite showing binary categories, these measurements are still relative and lose meaning when not compared to another group. The main limitation in this is there are no clear benchmarks to definitively say that a cell is clearly more button-like or zipper-like on its own. In addition, all assessments are image-based and focused on morphological differences. While junction morphology can provide insights into vascular outcomes, our study is limited by the lack of functional assessments such as transport-based vessel permeability studies [13,37] or transendothelial electrical resistance[7–12] to pair with morphological features. Finally, while the LEC-lined channels of microfluidic model we used approximates a cylindrical lymphatic vessel structure, direct comparisons to in vivo vasculature and related junction morphologies were not performed.

In summary, we presented a 3-step semi-automated image analysis pipeline that sufficiently sections LECs and quantifies LEC junction size and shape metrics to correlate with button- and zipper-like junction morphologies. This approach blended metrics previously used to describe endothelial junctions, such as junction thickness, number and size of junctions, with new metrics such as branchpoints and endpoints. Although radial intensity is not a new metric, our study leverages it in a new way to link VE-cadherin localization to junction morphology. No single image-derived metric fully defines a button-like or zipper-like junction, and PCA shows that multiple, complementary metrics can describe distinctions between morphologies. Junction and monolayer continuity can be described through branchpoints and endpoints, which can be reported individually but describe junctions most effectively when paired. Junction size and fragmentation is best defined by the relationship between object size and objects per cell, while localization of VE-cadherin across the cell body captures the dynamic nature of VE-cadherin expression. Overall, the analysis classified button-like junctions as being driven by metrics associated with fragmentation and discontinuities, while zipper-like junctions were dominated by VE-cadherin continuity and expression at the cell membrane. The insights gained from this study are bridging gaps for understanding the complexity of lymphatic architecture and LEC junction dynamics. While the current study uses TNF-α treatment with VE-cadherin response was used to validate this pipeline for LECs, the pipeline was designed to be adaptable and potential applications to other environment conditions and junction proteins in the future. Moreover, as research into lymphatic vasculature continues, it is important to develop robust tools for quantifying vessel features that can then be correlated to endothelial or vascular function.

## 5. Conclusions

In summary, we presented a 3-step semi-automated image analysis pipeline that sufficiently sections LECs and quantifies LEC junction size and shape metrics to correlate with button- and zipper-like junction morphologies. This approach blended metrics previously used to describe endothelial junctions, such as junction thickness, number and size of junctions, with new metrics such as branchpoints and endpoints. Although radial intensity is not a new metric, our study leverages it in a new way to link VE-cadherin localization to junction morphology. No single image-derived metric fully defines a button-like or zipper-like junction, and PCA shows that multiple, complementary metrics can describe distinctions between morphologies. Junction and monolayer continuity can be described through branchpoints and endpoints, which can be reported individually but describe junctions most effectively when paired. Junction size and fragmentation is best defined by the relationship between object size and objects per cell, while localization of VE-cadherin across the cell body captures the dynamic nature of VE-cadherin expression. Overall, the analysis classified button-like junctions as being driven by metrics associated with fragmentation and discontinuities, while zipper-like junctions were dominated by VE-cadherin continuity and expression at the cell membrane. The insights gained from this study are bridging gaps for understanding the complexity of lymphatic architecture and LEC junction dynamics. While the current study uses TNF-α treatment with VE-cadherin response was used to validate this pipeline for LECs, the pipeline was designed to be adaptable and potential applications to other environment conditions and junction proteins in the future. Moreover, as research into lymphatic vasculature continues, it is important to develop robust tools for quantifying vessel features that can then be correlated to endothelial or vascular function.

## Supplementary Materials

The following supporting information can be downloaded at: https://www.mdpi.com/article/doi/s1, Figure S1: title; Table S1: title; Video S1: title.

## Author Contributions

Conceptualization, S.M.L., A.K.F., and C.F.W.; data curation, S.M.L. and A.K.F.; formal analysis, S.M.L. and A.K.F.; funding acquisition – C.F.W; methodology, S.M.L., A.K.F.; project administration, C.F.W.; software, S.M.L. and A.K.F.; supervision, C.F.W.; validation, S.M.L.; visualization, S.M.L. and A.K.F.; writing—original draft, S.M.L. and C.F.W.; writing—review and editing, S.M.L. and C.F.W. All authors have read and agreed to the published version of the manuscript.

## Funding

This work was supported by the National Science Foundation Research Experience for Undergraduates (NSF REU) program (Grant EEC-2150076 to support A.K.F.), National Science Foundation Engineering Research Initiation Award (Grant 2138841 to C.F.W), and National Institutes of Health/National Cancer Institute R03 Small Grants Award (Grant 5R03CA267449-02 to C.F.W.).

## Data Availability Statement

The analysis pipeline information, and raw data supporting these conclusions is contained within the article or supplementary material

## Acknowledgments

The authors would like to thank Victoria Bicchieri and Dr. Samrat Dutta for their assistance and training for image acquisition, Lily Goudreau for her assistance with image analysis, and Dr Brian Ruliffson for foundational lymphatics work. During the preparation of this study, Copilot, MATLAB R2025a was used for the purposes of generating code for integration of CellPose, nuclei identification, exclusion criteria, and Dice Coefficient implementation in MATLAB script. The authors have reviewed and edited the output and take full responsibility for the content of this publication.

## Conflicts of Interest

The authors declare no conflict of interest.

## Appendix A Appendix A.1

**Table 1.** Descriptions of the metrics for quantifying zipper-like and button-like VE-cadherin junctions in LEC.

| Metric | Program | Definition | Significance | Illustration |
| --- | --- | --- | --- | --- |
| Radial Intensity | CellProfiler | Sections of cells radiate from the center of the cell to the membrane, each section measures average intensity in that region. | Radial intensity shows how proteins of interest localize in the cell.<br>Intensity of junction proteins shift from the outer cell membrane to internal structures as junctions begin to button. | Figure 4 |
| Skeletal Branchpoints | CellProfiler | Pixels in the foreground with greater than 2 pixels in the foreground. | Branchpoints occur when a line forks.<br>Jagged individual junction clusters lead to multiple small branching structures with buttoning. | Figure 5 |
| Skeletal End-points | CellProfiler | Pixels in the foreground of masked images with one neighboring foreground pixel. | Endpoints occur when a shape starts and ends in a point.<br>Endpoints increase in number as junctions fragment due to buttoning. | Figure 5 |
| Object Size | Fiji | Area of pixel groupings in the foreground of an image mask. | Shorter and discontinuous clusters of foreground pixels appear at the membrane leading to smaller junction fragments due to buttoning. | Figure 6 |
| Objects per Cell | Fiji | Number of whole pixel groupings in the foreground is divided by the number of identified cells in the image. | Multiple discontinuous junctions appear at the membrane and become separate foreground pixel clusters around a cell due to buttoning. | Figure 6 |
| Junction Thickness | Fiji | Widths of junctions averaged across the image. | Zipper-like, junctions tend to be long and thin structures.<br>Button-like junction disruptions are jagged with increased junction width. | Figure 7 |

**Table 2.** Description of how image data moves through the pipeline, where images originate from, and their purpose within the pipeline.

| Image | Input Program | Output Program | Description |
| --- | --- | --- | --- |
| RGB/Greyscale<br>VE-cadherin<br>channel | MATLAB,<br>CellProfiler | N/A | Images taken from microscope |
| RGB/Greyscale<br>Nuclear chan-<br>nel | MATLAB | N/A | Images taken from microscope |
| VE-cadherin<br>Border Mask | (CellProfiler) | MATLAB | Image mask outlining cell borders, can be used to verify cell sectioning |
| VE-cadherin<br>labels | CellProfiler | MATLAB | Object matrix from MATLAB, will not include any visuals, and will appear as a black image |
| DAPI Centroid<br>Mask | CellProfiler | MATLAB | Image mask of nuclei centroids |
| Thresholded<br>Mask | Fiji | CellProfiler | Image mask of a CellProfiler processed VE-cadherin image |
| Distance Map | N/A | Output (Fiji) | Heatmap of a thresholded mask image where thickness determines heatmap hue |

## Disclaimer/Publisher’s Note

The statements, opinions and data contained in all publications are solely those of the individual author(s) and contributor(s) and not of MDPI and/or the editor(s). MDPI and/or the editor(s) disclaim responsibility for any injury to people or property resulting from any ideas, methods, instructions or products referred to in the content.

## Notes

### Competing Interest Statement

The authors have declared no competing interest.

